# The spectral identity of foveal cones is preserved in hue perception

**DOI:** 10.1101/317750

**Authors:** Brian P. Schmidt, Alexandra E. Boehm, Katharina G. Foote, Austin Roorda

**Affiliations:** School of Optometry and Vision Science Graduate Group, University of California, Berkeley, USA, 94720

## Abstract

Organisms are faced with the challenge of making inferences about the physical world from incomplete incoming sensory information. One strategy to combat ambiguity in this process is to combine new information with prior experiences. We investigated the strategy of combining these information sources in color vision. Single cones in human subjects were stimulated and the associated percepts were recorded. Subjects rated each flash for brightness, hue and saturation. Brightness ratings were proportional to stimulus intensity. Saturation was independent of intensity, but varied between cones. Hue, in contrast, was assigned in a stereotyped manner that was predicted by cone type. These experiments revealed that, near the fovea, long (L) and middle (M) wavelength sensitive cones produce sensations that can be reliably distinguished on the basis of hue, but not saturation or brightness. Taken together, these observations implicate the high-resolution, color-opponent parvocellular pathway in this low-level visual task.

## Introduction

Incoming sensory information is inherently noisy and ambiguous. The retinal image and the subsequent signals encoded by the photoreceptors can be interpreted infinitely many ways. Helmholtz (1924) posited that perception represents the brain’s best guess about the state of the world after taking into account both the ambiguous incoming signals and prior experience. Investigating the rules through which incoming sensory signals are combined with prior evidence is an important area of brain research (Knill and Pouget, 2004). Here, we studied the color appearance of light targeted to a single receptor in order to elucidate the rules the visual system follows when presented with impoverished information from its primary sensory neurons. Understanding these rules will provide insight into how the visual system handles uncertainty in more naturalistic tasks as well.

A light of fixed wavelength that is sufficiently small in diameter and weak in intensity will fluctuate in appearance across presentations (Krauskopf, 1964; Krauskopf and Srebro, 1965; Otake and Cicerone, 2000). To understand why, consider a spot small enough to stimulate only a single cone. An individual cone is colorblind; information about wavelength is not retained after a photoreceptor captures a photon (Rushton, 1972). The visual system computes color information by comparing relative activity across the three cone types. If light falls on only a single cone, the visual system will be missing samples from the other two cone types and the color of the spot will be ambiguous. One possible solution to this problem is that the visual system could use prior experience to infer the activity of the two missing classes when judging the color of the spot (Brainard et al., 2008). The reason why the spot fluctuates in appearance from trial to trial is that the eye makes tiny movements from moment to moment. As a result of this incessant movement, the light will strike a different cone each time it is flashed on the retina and, thus, a small spot of light with a fixed wavelength will fluctuate in appearance across presentations (Krauskopf, 1964; Krauskopf and Srebro, 1965; Krauskopf, 1978).

In a pioneering examination into this phenomenon, Krauskopf and Srebro (1965) asked subjects to match small, dim flashes on a dark background to monochromatic light. Under these conditions, they discovered that matches fell into one of two distinct clusters: one centered around orangish-red wavelengths and the other blueish-green. They hypothesized that (1) the two perceptual distributions corresponded to two distinct detectors – L- and M-cones, respectively – and that (2) white sensations arose when an L- and an M-cone were activated together. Due to technological limitations, they were unable to isolate single cones, identify the cone type targeted or repeatedly target the same cone. Thus, a conclusive test their hypotheses was not possible. More recently, Hofer et al. (2005b) improved single cone isolation with adaptive optics, which corrects for each subject’s optical imperfections. Contradicting the second hypothesis of Krauskopf and Srebro (1965), they concluded that the frequency of white reports was too high to be caused exclusively by trials where L- and M-cones were stimulated together: at least some single cone flashes produced white sensations. The authors also argued that the variability they observed was too great for the response of a single cone to depend only on its spectral class. However, Hofer et al. (2005b) were not able to target light to receptors of known spectral type and, therefore, could not directly relate cone activity to color reports.

Recently, we combined high-resolution eye-tracking with adaptive optics to additionally compensate for natural eye movements (Arathorn et al., 2007; Harmening et al., 2014). With this technology, we stimulated individual cones and identified their spectral type (Sabesan et al., 2015). Our results confirmed the prediction that most of the variability in chromatic percepts could be attributed to the type of cone that was targeted (Sabesan et al., 2016; Schmidt et al., 2018). On average, L-cones mediated red sensations, while M-cone trials gave rise to green or blue, depending on the background context. At a supra-threshold intensity and against a white background (Sabesan et al., 2016), a majority of trials were judged white. This was consistent with the observations of Hofer et al. (2005b) and contradicted the second hypothesis of Krauskopf and Srebro (1965) that light absorbed by a single cone always produces saturated color percepts. However, unlike the older studies, our experiments were conducted on a white background which may reduce variability in color ratings (Schmidt et al., 2018; Hofer et al., 2012).

The discrepancy between these studies raises new questions about how the visual system parses hue and achromatic sensations from a mosaic of detectors that are individually colorblind. Firstly, do sensations from individually targeted L and M cones truly vary in saturation (amount of whiteness)? More specifically, do single-cone color percepts exist on a continuum, with each cone producing a mixture of hue and achromatic sensation (Wool et al., 2018) or is there a discrete subclass of cones wired specifically into chromatic or achromatic pathways (Neitz and Neitz, 2017)? In our prior work, subjects reported on their perception with only a single color name (Sabesan et al., 2016; Schmidt et al., 2018). The limited range of response options may have obscured more subtle variation in hue and saturation. Secondly, color and brightness perception of small spots are known to change with intensity (Kaiser, 1968; Weitzman and Kinney, 1969). However, the mechanism underlying this phenomenon is unclear. A higher luminance spot will both activate more cones and do so more strongly. We sought to understand whether individually targeted cones would systematically change in appearance – perhaps from white to colored – as the number of photons per flash was increased.

The relationship between stimulus intensity and color appearance at a cellular-scale was quantified using precise measurements of the sensation elicited by each spot. The results revealed that subjects used color terms in a stereotypical manner predicted by cone type, but largely independent of stimulus intensity.

## Methods

### Subjects

Three highly experienced subjects (two male, one female [S20092]) between the ages of 27 and 34 participated in the study. All subjects had normal color vision and two (S20076 and S20092) were authors of the study. All procedures were approved by the Institutional Review Board at the University of California Berkeley and adhere to the tenets of the Declaration of Helsinki. Informed consent was obtained from each subject before the experiments. At the start of each session, cycloplegia and mydriasis were induced with drops of 1.0% tropicamide and 2.5% phenylephrine hydrochloride ophthalmic solution.

### Cone-resolved imaging and stimulation

A multi-wavelength adaptive optics scanning laser ophthalmoscope (AOSLO) was used to image and present stimuli to the retina. The objective of this study was to confine a small stimulus probe (543 nm) to targeted locations on the retina, *i.e*. individual cones. Monochromatic imperfections were measured with a wavefront sensor (940 nm) and corrected with a deformable mirror (Roorda et al., 2002). Imaging was performed with 840 nm light, which was collected into a photo-multiplier tube through a confocal pinhole and rendered into a video stream. The video stream was registered to a reference image in real-time with a strip based cross-correlation procedure (Arathorn et al., 2007). The output of this procedure produced retinal coordinates that were used to drive an acousto-optic modulator, a high-speed optical switch, which delivered 543 nm visual stimuli to the retina whenever the raster scan passed over the targeted cell. Chromatic aberration was measured and corrected with established procedures (Harmening et al., 2012). In these experiments, a 512 × 512 pixel imaging raster subtended a 0.95° field, with a sampling resolution of ~0.11 arcmin pixel^−1^.

The challenges involved in targeting single cones and the technology for overcoming these challenges has been described elsewhere (Roorda et al., 2002; Arathorn et al., 2007; Harmening et al., 2012, 2014; Sincich et al., 2015) and a full consideration of the issues involved in stimulating individual receptors is beyond the scope of this paper. However, before analyzing the psychophysical results, the ability of our system to confine stimulus light to the targeted cone is worth considering. Briefly, there are three main sources of noise that causes a point source to be blurred at the retinal plane: eye-motion, residual optical imperfections and forward light-scatter. All three potentially limit image quality and the isolation of single cones. Forward-scatter manifests as a large, dim halo surrounding the peak of the point-spread-function. The magnitude of scatter relative to the core of the PSF has been estimated to be 1:10,000 (Hofer et al., 2005b; Harmening et al., 2014; van den Berg et al., 2010). In the dark, these scattered photons may have visual consequences. However, in the present work, a photopic background (see below) raised thresholds for each cone and minimized the influence of uncontrolled scatter.

To assess the impact of any residual blur or eye-motion, the light profile of the stimulus was modeled at the retinal plane and the fraction of light absorbed by the cone mosaic was computed (Harmening et al., 2014). The location of each stimulus was first determined. During each trial of the experiment, a video of the retina was recorded and the location of the stimulus in each frame was indicated with a digital mark. The digital mark was recovered from each frame after the experiment to assess how closely the actual stimulus was delivered to the desired location. The contours in the left plot of Figure 1A represent the distribution of stimulus delivery locations over all frames from all trials during an example session. The highest density of those distributions is concentrated at the center of the five targeted cones. Next, the image of each stimulus on the retina was modeled by convolving a near diffraction-limited (0.05 D residual defocus) point spread function with the 3 × 3 pixel (~0.35 arcmin) stimulus; 0.05 D was chosen as a conservative magnitude of uncorrected optical aberration (Harmening et al., 2014). The contours in right panel of Figure 1A demonstrate that most of the stimulus light was confined to the targeted cone, even after accounting for uncorrected eye-movement and optical defocus. At the eccentricity (1.5°) tested in this subject (S20076), cones were separated by approximately 9-12 pixels or slightly more than one arcmin. Had cones been more tightly packed, for instance closer to the fovea, a greater fraction of the light would have been inadvertently delivered to neighboring cones.

**Figure 1:**
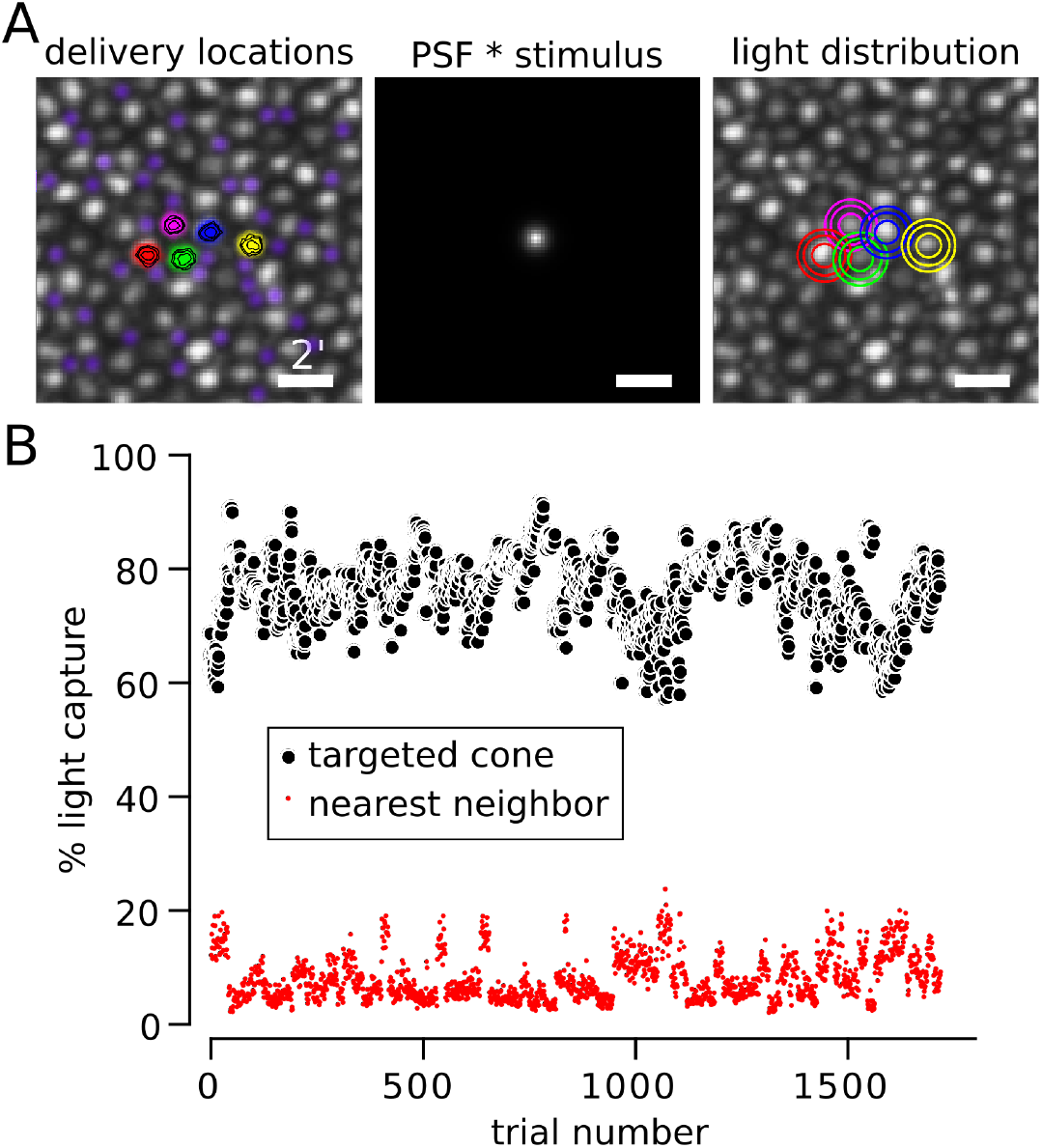
Targeting light to individual cones. **A**. *Left*: Delivery locations of 5 cones from S20076. The location of the stimulus was recovered on each frame of each trial (15 frames, 500 ms) and recorded. Contours indicate that delivery locations were concentrated at cone centers. Rods were pseudo-colored purple to distinguish them from cones (the larger, gray-scale, cells). *Middle*: 3 × 3 pixel stimulus convolved with a near diffraction-limited PSF (6.5 mm pupil with 0.05 diopters of defocus (Harmening et al., 2014)). *Right*: density profile of light capture in each cone computed by summing the PSF ∗ stimulus at each delivery location. For both *Left* and *Right* plots contours encompass 50, 80 and 90% of delivered light from smallest to largest. Scale bar = 2 arcmin. **B**. Estimated percentage of light captured by the targeted cone (black circles) and its nearest neighbor (red circles) during each trial. Light spread was modeled as described in **A** and each cone aperture was assumed to be Gaussian with a width at half height of 0.48 relative to the diameter of cone inner segments (MacLeod et al., 1992; Chen et al., 1993). Trials with delivery errors greater than 0.35 arcmin were excluded from analysis.

In Figure 1B, we modeled how much light was absorbed by the targeted cone versus its nearest neighbor after taking into account the aperture of cones at this eccentricity (Harmening et al., 2014), which was assumed to be a Gaussian profile with a full-width half-max of 0.48 of the cone inner segment diameter (MacLeod et al., 1992; Chen et al., 1993). This analysis was repeated for each trial. Of the light absorbed by the mosaic, the targeted cone captured, on average, 76.4% (*σ* = 6.2%) in this subject. The next nearest neighbor captured 8.1% (*σ* = 4.0%), while most of the remaining 15.5% of absorbed light fell on the five other adjacent cones. Therefore, in theory (Harmening et al., 2014), the targeted cone in this subject absorbed 9-10 times more light than any of its neighbors. Allowing the cone aperture to vary over physiologically plausible values (0.4-0.56) (MacLeod et al., 1992; Chen et al., 1993) caused only modest impact on the estimated percentage of light captured by the targeted cone (80.2 - 72.1). This analysis produced similar estimates for the other two subjects (aperture=0.48; S20053: targeted cone *μ* = 67.3%, *σ* = 6.6%, nearest neighbor *μ* = 9.9%, *σ* = 3.6%; S20092: targeted cone *μ* = 82.1%, *σ* = 4.2%, nearest neighbor *μ* = 6.7%, *σ* = 2.9%. *μ* = mean, *σ* = standard deviation). The lower light capture in S20053 was predominantly driven by a higher packing density of cones at the location tested (~1°). The potential influence of greater light capture by neighboring cones is considered in the Results section.

### Stimulus and background parameters

Cones were targeted with spots (543 nm; 500 ms; 0.35 arcmin) that varied in intensity. Flash intensity was defined in linearized arbitrary units (a.u.) of the maximum intensity presented. A flash strength with a.u. = 1 delivered approximately 3.69 × 10^6^ photons to the cornea or 5.19 log_10_ Trolands (Nygaard and Schuchard, 2001). The background in these experiments was composed of four sources: (1) an invisible 940 nm wavefront sensing beam, (2) a dimly visible 840 nm imaging raster (2.02 log_10_ Trolands), (3) leak from 543 nm stimulation channel (1.99 log_10_ Trolands) and (4) an external projector. The external projector was imaged onto the subject’s retina in Maxwellian view. Before each session, the subject adjusted the chromaticity and luminance of the projector until the entire mixture appeared white. The luminance was approximately 2.52 log_10_ Trolands. Together, the four background sources produced ~2.73 log_10_ Trolands. Thus, at one a.u. the stimulus was approximately 290 times more luminous than the background. Each cone location was tested at three intensities. Flash strength was chosen to sample the entire range of the frequency of seeing curve. S20053 and S20076 were tested at identical stimulus intensities. S20092 was first tested at the intensities used for the other two subjects. However, due to overall lower sensitivity to the stimulus, S20092 was subsequently tested at slightly higher intensities.

### Psychophysical procedure

At the start of each psychophysics session, a high SNR image was collected from an average of 90 frames. From that image, the locations of four to six cones were marked. Each cone was targeted ten times at each of three intensities and an additional 10% of blank trials were added. Trials were randomly interleaved between cone locations and stimulus intensities. A dataset of roughly 40-100 cones from each subject was collected over a minimum of ten sessions, which were spread over multiple days. Where possible, cones were targeted contiguous to previously tested locations. The analyses presented here consider all of the cones tested in each subject.

The subject initiated each trial with a button press, which was accompanied by an audible beep. After each flash, the subject rated the brightness of each stimulus on a scale from 0 to 5 (brightest). The subjects were given at least three practice sessions (~500 trials) to develop an internal criterion for assigning ratings. No reference stimulus or feedback was given. Trials that received brightness ratings above zero were also rated for hue and saturation (Gordon et al., 1994). The subject indicated the percent of red, green, blue, yellow and white contained in each stimulus using five button presses such that each press represented 20% (5×20%=100%). At least three full practice sessions were completed to develop familiarity with the task and the range of percepts experienced.

### Cone classification

A small mosaic of cones 1-2° from the fovea in each subject was selected for study. The spectral class of targeted cones were identified using densitometry (Hofer et al., 2005a; Roorda and Williams, 1999; Sabesan et al., 2015). Densitometry measurements were collected in imaging sessions separate from the psychophysical experiments. In one subject (S20092), we were unable to collect densitometry data due to time limitations.

### Analysis

Data were aggregated across all experimental sessions. Responses from each trial were organized by cone location and stimulus intensity. Before inclusion into the data set, each trial was analyzed for delivery error. The location of the stimulus on each frame of the recorded video was recovered. Delivery error was defined as the standard deviation of those recovered locations. Trials with delivery errors greater than 0.35 arcmin were discarded.

Frequency of seeing (FoS) curves were computed by binarizing brightness ratings: ratings above one were seen. FoS data was analyzed on both a cone-by-cone basis as well as in aggregate over all cones within a single class. In both cases, the data were fit with a Weibull function, Φ, defined as:

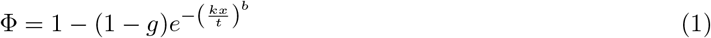

where

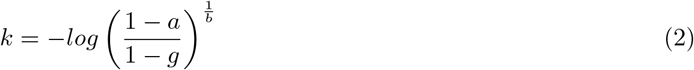

In this parameterization of the Weibull function, *g* represented the performance measured during blank trials, *t* was the threshold and *a* was the proportion correct that defined the threshold (here *a* = 0.5). The slope of the function was defined by *b*. Model parameters were fit to the data using a maximum likelihood routine. Only *t* and *b* were treated as free parameters.

Hue and saturation were analyzed for all seen trials. Responses from each trial were converted into a uniform appearance diagram (UAD) (Gordon et al., 1994; Abramov et al., 2009). The axes in this space were defined as 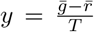 and 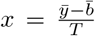, where *ḡ*, 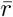, *ȳ*, 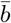 correspond to the number of green, red, yellow, blue responses, respectively, and *T* was the total number of button presses, which was five. Saturation was computed as distance from the origin in city block metric (Gordon et al., 1994), *i.e* |*x*| + |*y*|. A purely white response falls at the origin of this space, while a purely saturated response falls along boundary where |*x*| + |*y*| = 1. Hue angles relative to the origin were also computed from the *x* and *y* position of each data point. *x* > 0 and *y* = 0 represented an angle of 0°. Trials with pure white responses (5 white button presses) were excluded from this analysis because the angle is undefined.

The percent variance in hue angle, *θ*, or saturation, **S**, explained by cone type was computed following a procedure adapted from Carandini et al. (1997). The mean square difference between two sets of responses **x** = *x_c_* and **y** = *y_c_* was computed:

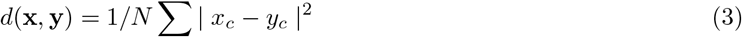

where the sum was taken over tested cones, *c*, and *N* was the number of tested cones. The percent variance explained, **% variance**, by cone type was then expressed as:

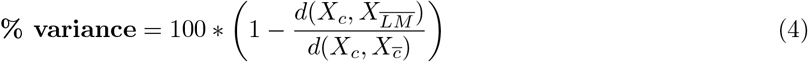

where *X_c_* was the mean hue angle or saturation for each cone, 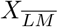 was the mean hue angle or saturation across L- and M-cones separately and 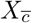 was the mean hue angle or saturation across all cones, regardless of cone type.

All analyses were carried out in MATLAB. Analysis code and raw data may be freely downloaded from GitHub (https://github.com/bps10/SchmidtBoehmFooteRoorda_2018).

## Results

Cone photoreceptors in three human volunteers were imaged and stimulated with an adaptive optics scanning laser ophthalmoscope (AOSLO). Figure 2 illustrates the results from an example session. Five cones were selected: one M-cone and four L-cones. Each cone was tested ten times at three different stimulus intensities and an additional 10% of trials were blanks. Stimulus intensity was randomized across trials. After each flash, the subject first judged the brightness on a scale from 0 (not seen) to 5. Each subject developed his or her own criterion for brightness during practice sessions. Secondly, on trials that were seen the subject additionally judged the hue and saturation of each flash with a rating scale (Abramov et al., 2009; Gordon et al., 1994). The subject indicated the proportion of red, green, blue, yellow and white in increments of 20% (for a total of 100%). For instance, a desaturated teal might be 20% blue, 20% green and 60% white. Hue scaling data were then transformed into a uniform appearance diagram (UAD) (Figure 2D). In a UAD the x-axis indicates to the strength of yellow-blue sensations 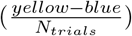 and the y-axis indicates green-red bias 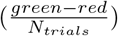. White falls at the origin of this diagram and completely saturated responses (0% white) fall on the outer edge, as defined by the dotted lines. Saturation was computed as the distance from the origin (in city-block metric).

**Figure 2:**
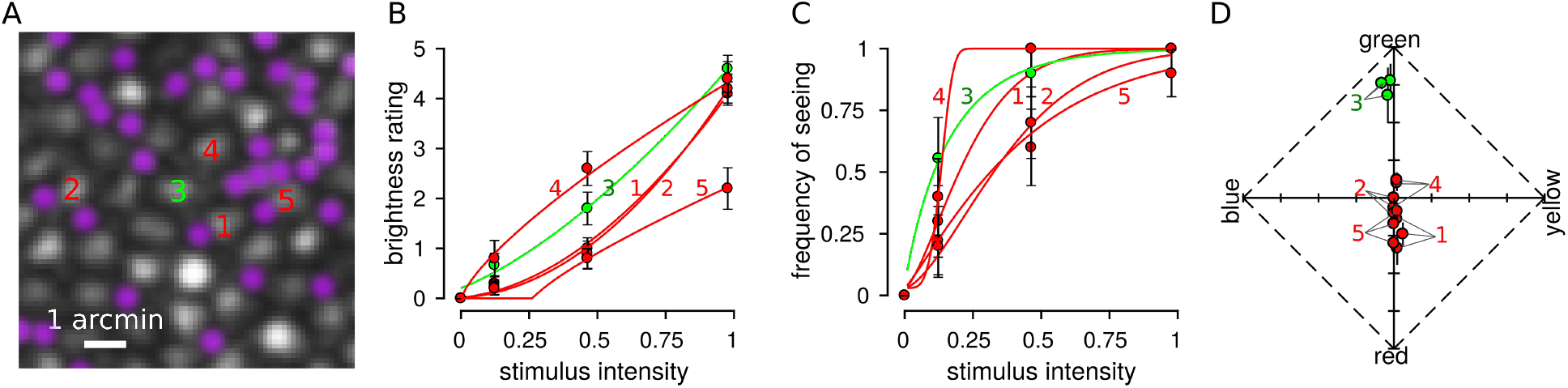
Hue, saturation and brightness scaling from spots targeted to single cones. **A**. AOSLO image from S20076 at 1.5° eccentricity. Numbers indicate the five cone centers that were selected for targeting during the experimental session. A roughly equal proportion of rods and cones are visible at this eccentricity (rods have been pseudo-colored purple). Scale bar indicates 1 arcmin. **B**. On each trial, a flash was targeted at one of the selected location and trials were randomly interleaved between the five locations. After each flash, the subject rated brightness on a scale from 0 to 5 and data were fit according to Steven’s Law (Eq. 5). A brightness rating greater than zero indicated the trial was seen. **C**. FoS data were fit with Weibull functions (Eq. 1). **D**. Subjects additionally reported the hue and saturation of the flash with a scaling procedure. The results from hue and saturation scaling from the five tested cones are plotted in a uniform appearance diagram (Abramov et al., 2009; Gordon et al., 1994). Each cone was tested at three intensities and the average response across all seen trials is plotted for each intensity. Error bars represent SEM; some error bars are smaller than the size of the symbol. Colors denote cone type (green = M-cones, red = L-cones.)

The cone mosaics and position of targeted cones from three subjects are plotted in Figure 3. These datasets were amassed over numerous experimental sessions. Targeted cone locations are indicated by the presence of a pie chart. The pie chart represents a histogram of all button presses (red, green, blue, yellow, white) from all seen trials. The tested locations were between 1 and 2 degrees of eccentricity. The region targeted in S20053 was closest to the foveal center (~1 °) and had the highest cone density. Subjects S20053 and S20076 used “white” more frequently than any other color category. Subject S20092 used hue categories more often than “white.” These plots illustrate that hue and saturation judgments were variable from cone to cone and possibly even between those with the same photopigment. Below, we examine these observations in more detail.

**Figure 3:**
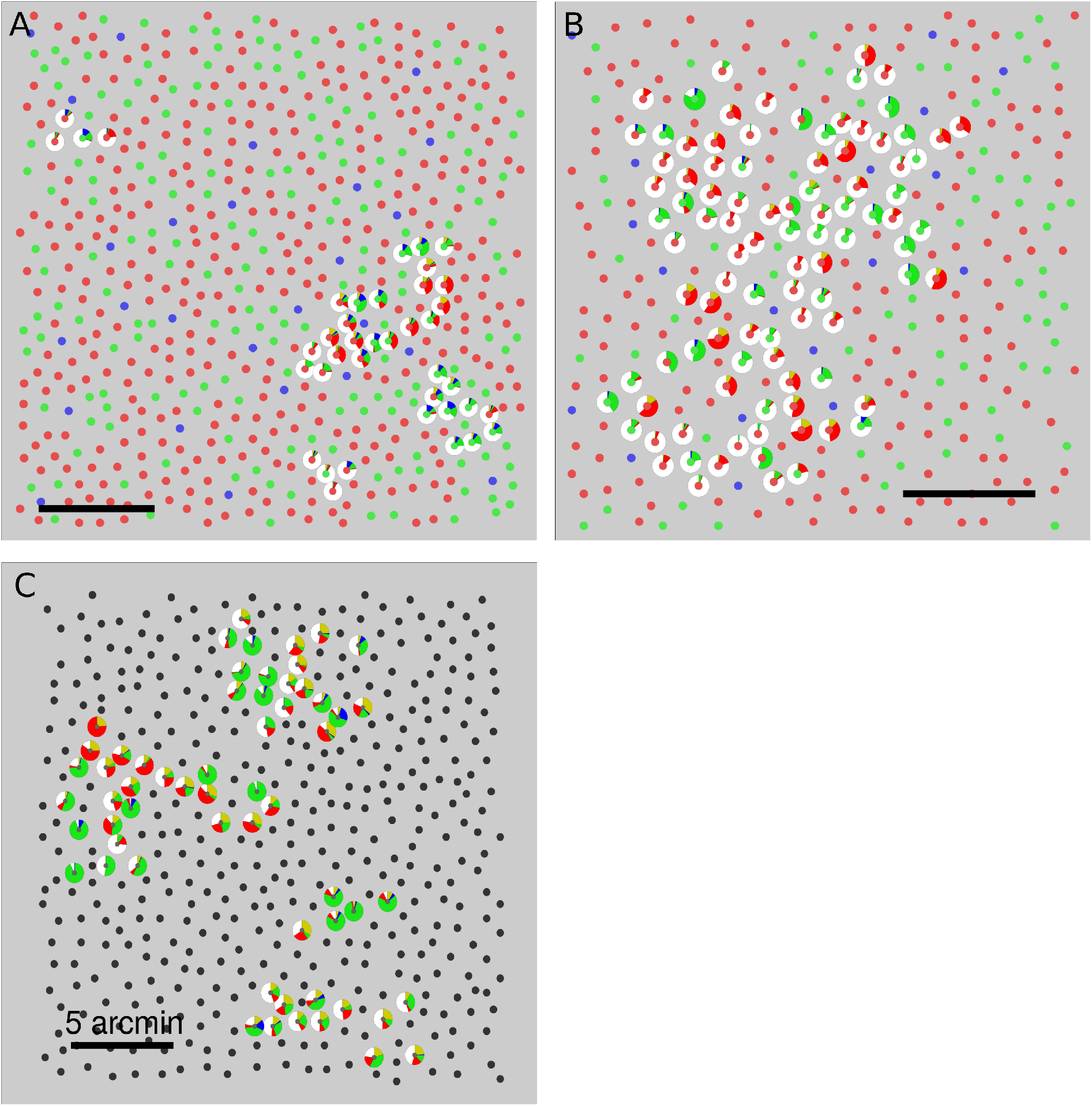
Spatial variation of hue and saturation scaling. Targeted cones are surrounded by a pie chart that represents the fraction of times a given color was reported. Each seen flash was scaled with five button presses. Pie charts illustrate all button presses from every seen trial across all three stimulus intensities. Cone locations are denoted by smaller circles. The spectral class of each cone is indicated by the color. L-cones = red, M-cones = green, S-cones = blue, Unclassified = black. **A**. S20053, **B** S20076, **C**. S20092. Scale bars indicate 5 arcmin or ~24*μ*m.

### Influence of stimulus intensity on detection and brightness

The influence of stimulus intensity on detection and brightness judgments were first analyzed across cone classes. Trials were grouped according to cone type and stimulus intensity. FoS was then computed from binarized brightness ratings (ratings above 0 were seen). Figure 4 reports the FoS across our three subjects. In S20053 and S20076, L- and M-cone thresholds (defined here as 50% FoS) were similar (Table 1). This finding is expected based on the sensitivity of L- and M-cones to our 543 nm stimulus (Stockman and Sharpe, 2000). Single cone thresholds were higher on average in S20092, but within the normal range expected for healthy volunteers (Harmening et al., 2014; Bruce et al., 2015).

**Figure 4:**
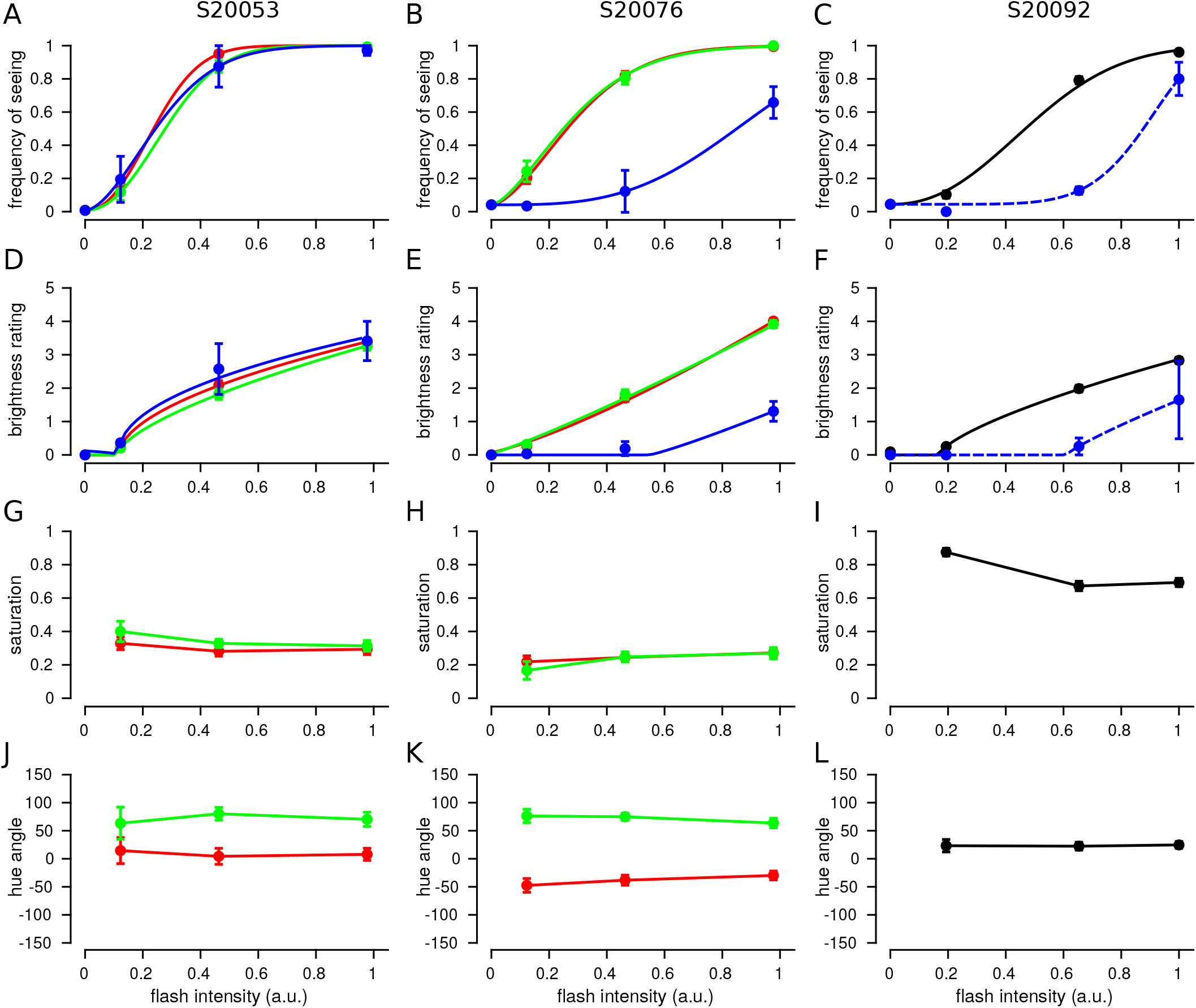
Brightness is dependent upon stimulus intensity, but hue and saturation are not. Trials from each cone were grouped according to flash intensity and the mean response was computed. The data plotted represent the mean and SEM across tested cones. Each column represents data from one subject. *Left*: S20053, *middle*: S20076, *right*: S20092. **A-C**: Frequency of seeing. Data were best fit with Weibull functions (Eq 1). **D-F**: Mean brightness ratings for each subject. Data were fit with exponential Steven’s Law functions (Eq 5). **G-I**: Mean saturation ratings. **J-L**: Mean hue angles. Color denotes spectral identity of cone targeted: red = L, green = M, blue = S, black = unclassified L or M cone, blue dotted = purported S-cones. *Left column*: S20053. Error bars represent SEM.

**Table 1:** FoS psychometric function fits

|  | S20053 |  |  |  | S20076 |  |  |  | S20092 |  |  |  |
| --- | --- | --- | --- | --- | --- | --- | --- | --- | --- | --- | --- | --- |
| | N | $t$ | $b$ | log-likelihood | N | $t$ | $b$ | log-likelihood | N | $t$ | $b$ | log-likelihood |
| L-cones | 22 | 0.237 | 2.206 | 15.74 | 77 | 0.263 | 1.656 | 76.84 | 60 | 0.468 | 2.25 | 70.52 |
| M-cones | 18 | 0.278 | 2.161 | 15.84 | 33 | 0.25 | 1.531 | 39.37 |  |  |  |  |
| S-cones | 2 | 0.244 | 1.729 | 2.08 | 9 | 0.851 | 3.305 | 11.89 | 2 | 0.877 | 6.701 | 1.94 |

Data were fit according to the Weibull function defined in Eq 1. $t$ represents the threshold for 50% frequency of seeing in arbitrary stimulus units. $b$ controls the slope of the function.

Threshold for seeing was higher on trials targeted at S-cones. 543 nm light is about 300 times less effective at activating S-cones relative to either L- or M-cones (Stockman and Sharpe, 2000). Therefore, trials targeted at S-cones should be undetectable. In cases where a spot was detected, we assume that either neighboring L/M-cones absorbed some fraction of the light or the cone was mis-classified and was in reality an L- or M-cone. Threshold measurements have been previously used to elucidate S-cone topography near the fovea (Williams et al., 1981). The FoS curves measured in S20076 and S20092 approximately adhere to these expectations. In S20076, S-cone thresholds were significantly elevated relative to L- and M-cones. In S20092, two (out of 62) cones had elevated thresholds and were purported to be S-cones (Table 1). Only at the highest intensities tested did S-cone FoS increase above 50% in either subject (Figure 4B,C). In contrast, S20053 did not exhibit this behavior. S-cones thresholds were indistinguishable from L- and M-cones. This finding may indicate that single cones were less well isolated in this subject. However, were this the case, FoS should nonetheless be systematically lower than L/M-cones due to at least a fraction of the light falling on the targeted S-cone. A more likely explanation is the tested S-cones were mis-classified and were actually L- or M-cones. We consider these two possibilities below.

Brightness ratings from L-, M- and S-cones are displayed in Figure 4D-F. The three subjects in our study exhibited similar gross reports of brightness ratings, which increased predictably with stimulus intensity. The dependence of intensity *I* on perceived brightness, *ψ*, was modeled according to Steven’s Law (Stevens, 1966, 1961):

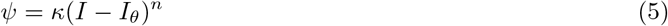

where *κ* represents a scaling constant and *I_θ_* has been interpreted as a threshold (Stevens, 1961). When *n* < 1 the relationship follows a compressive non-linearity and when *n* > 1 the non-linearity becomes expansive. In S20053 and S20092, *n* was less than one. This finding was consistent with a compressive non-linearity that has been previously reported for brightness scaling of small spots (Stevens, 1966, 1960). S20076 was best fit by a nearly linear relationship (Figure 4E; L-cones: *n* = 1.26; M-cones: *n* = 1.11).

S-cone targeted brightness ratings were expected to be lower than L/M-cone trials due to lower sensitivity to the stimulus wavelength. Judgments from S20076 and S20092 followed this expectation (Figure 4E&F). For the two S-cones targeted in S20053, brightness ratings were no different from L/M-cone trials. This observation is more consistent with the interpretation that the two S-cones tested in this subject were mis-classified by the densitometry procedure and were actually L- or M-cones. Of the three cone types, S-cone classification is the least reliable (Roorda and Williams, 1999; Hofer et al., 2005a; Sabesan et al., 2015). Weakly reflective L- and M-cones can, in some cases, be classified as S-cones.

### Hue and saturation do not depend on intensity

The results above demonstrated that FoS and brightness ratings depended on stimulus intensity. We next asked to what degree hue and saturation ratings correlated with stimulus intensity. In comparison to brightness, stimulus intensity did not substantially influence hue or saturation ratings over the range studied (Figure 4G-L). The largest shifts were observed between the lowest and highest energy stimuli. However, the standard error of the mean was high in low intensity conditions due to low FoS (< 0.25 in all subjects).

The influence of stimulus intensity was also analyzed on a cone-by-cone basis. In all three subjects, hue and saturation judgments were correlated across the two highest intensities (all comparisons, all subjects: *p* < 0.001) and had slopes close to unity. Figure 5 shows the results from S20076 (*hue*: *R*^2^ = 0.881, slope = 0.85; *saturation*: *R*^2^ = 0.514, slope = 0.81). The results from S20053 (*hue*: *R*^2^ = 0.735, slope = 0.76; *saturation*: *R*^2^ = 0.572, slope = 0.88) and S20092 (*hue*: *R*^2^ = 0.687, slope = 0.74; *saturation*: *R*^2^ = 0.503, slope = 0.63) were similar. Hue angles from cones with low saturation ratings (< 0.1) were excluded from the analysis (29 of 187 cones). Hue angle values below 0.1 were inherently noisy due to the small number of button presses that indicated the presence of a hue. Inclusion of low saturation cones did not materially change the results. Overall, these findings lend further support to the conclusion that hue and saturation judgments were largely independent of intensity.

**Figure 5:**
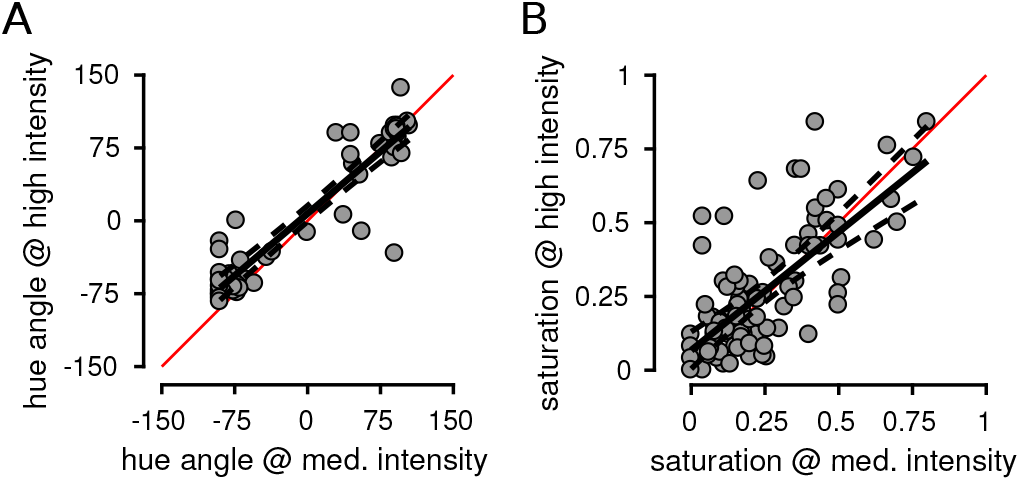
Hue and saturation judgments were correlated across stimulus intensities. Medium and high stimulus intensities were compared. Each point represents the mean response from an individual L- or M-cone across all seen trials at the indicated stimulus intensity. Data are shown from S20076. Hue angles (**A**) and saturation values (**B**) were highly correlated across stimulus intensity levels. Solid black line indicates best fit regression and dotted lines are 95% confidence intervals. Red line = unity.

### Variability in color sensations from cones with the same photopigment

Unlike stimulus intensity, cone type imparted a substantial bias on hue reports. Figure 4J-L reveals clear differences in the mean hue angle, *μ_θ_*, recorded from spots targeted at L- (S20076: *μ_θ_* = −35.6 ± 2.6°, *N_trials_* = 793; S20053: *μ_θ_* = 1.9±5.3°, *N_trials_* = 326) versus M-cones (S20076: *μ_θ_* = 74.5±2.7°, *N_trials_* = 421; S20053: *μ_θ_* = 80.5± 4.7°, *N_trials_* = 263). On the other hand, saturation was not dependent on spectral class (Figure 4G&H).

To understand this observation in more detail, color reports were analyzed on a cone-by-cone basis. Responses from each trial were grouped by the cone targeted, regardless of stimulus intensity. Figure 6 represents the data collected from each L- and M-cone in a UAD. The appearance of spots directed to L-cones clearly display a tendency towards reddish-yellow, while M-cone targeted spots were identified as green or greenish-blue in appearance. To quantitatively measure this tendency, the percentage of hue scaling variance explained by cone type was computed according to Eq. 4. Cone type accounted for 29.9% (N=40) and 41.4% (N=97) of the between cone variance in hue angle judgments in S20053 and S20076, respectively. Those numbers increased when cones with saturation values < 0.1 were excluded from analysis (S20053: 32.7%, N=38; S20076: 50.1%, N=83). In comparison, between cone variability in saturation was not predicted by cone type (S20053: 2.0%, N=40; S20076: 0.01%, N=97).

**Figure 6:**
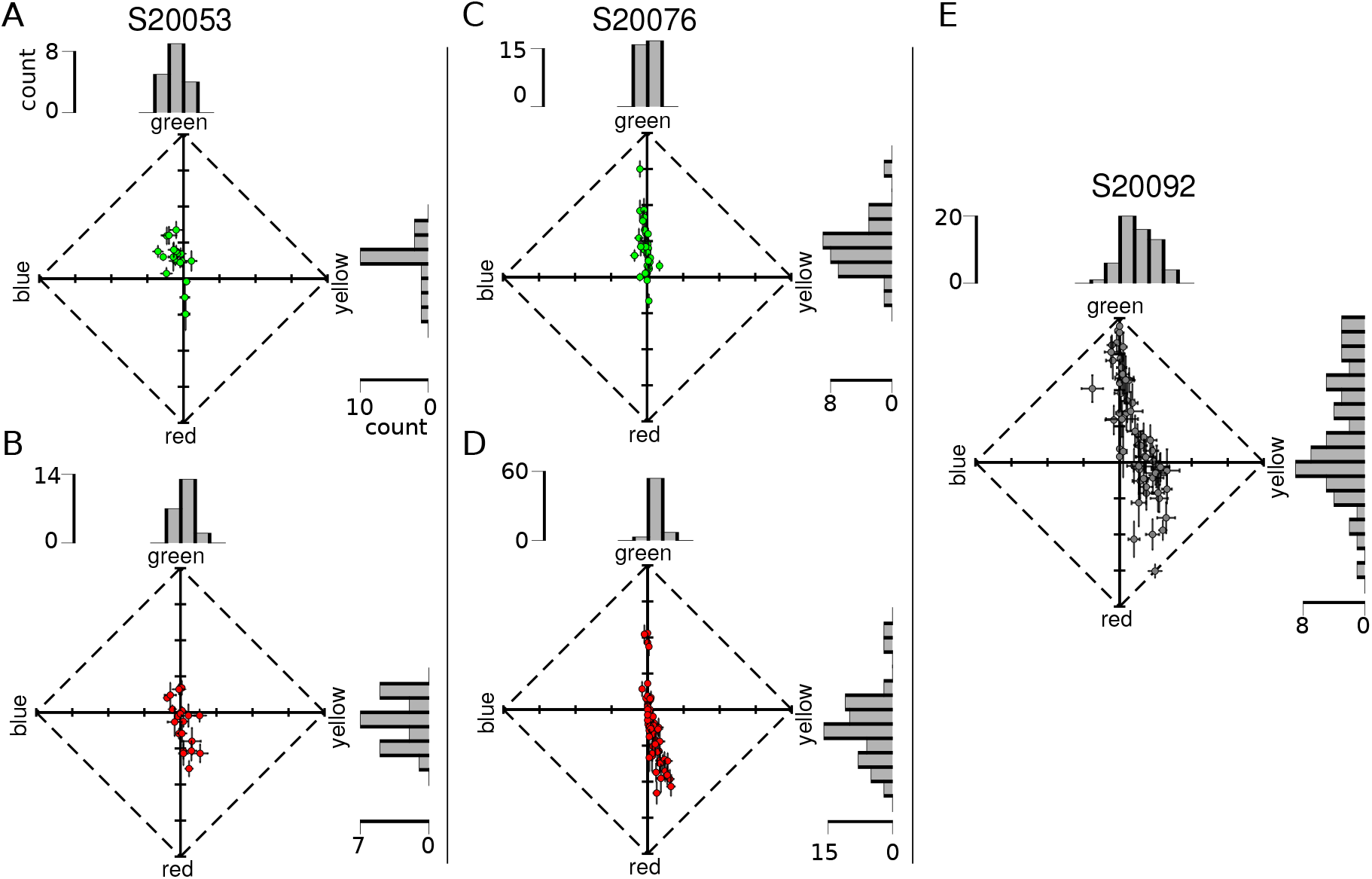
Hue and saturation reports from targeted cones. Each dot represents the mean response (of all seen trials, across all stimulus intensities) from an individual cone. Data are represented in a uniform appearance diagram, where the y-axis denotes a bias of responses towards greenness versus redness and the x-axis denotes a bias towards blueness versus yellowness. (**A**) M-cones and (**B**) L-cones tested in S20053. (**C**) M-cones and (**D**) L-cones tested in S20076. (**E**) Data from unclassified L- and M-cones in S20092. Error bars indicate SEM. Histograms above and to the right of each plot illustrate the distribution of responses along each dimension. Scale bars denote the number of cones in each bin.

Next, we asked whether cones sharing the same photopigment produced statistically distinguishable responses. As reviewed in the Introduction, Krauskopf and Srebro (1965) proposed that L-cones stimulated in isolation would produce red/yellow sensations, while M-cones would elicit green/blue reports. To address this hypothesis, we first ran a one-way analysis of variance (ANOVA) to establish whether hue/saturation responses differed significantly between cones (Figure 6). The *y-b* and *g-r* dimensions of UAD space were assessed separately. In all three subjects, there was a main effect for cone targeted in both response dimensions (*p* < 0.01 for all comparisons; S20053 *y-b*: F_39,766_=4.0, *g-r*: F_39,766_=10.6; S20076 *y-b*: F_96,2286_=8.3, *g-r*: F_96,2286_=23.8; S20092 *y-b*: F_59,1054_=6.5, *g-r*: F_59,1054_=13.0). Subsequently, post-hoc analyses (Tukey-Kramer) were run to determine which cones differed from one another.

The heatmaps in Figure 7 displays the statistical significance (log_10_ transformed) of each comparison. Each square represents a single post-hoc comparison between two cones. Results were organized according to cone class. Yellower squares indicate that mean responses did not differ; bluer squares indicate that the two cones elicited significantly different responses. If cones with the same photopigment produce a single color sensation, we should find a clump of yellow squares when cones of the same photopigment were compared and blue clumps corresponding to L- and M-cone comparisons. The results did not support the strongest form of this hypothesis. The heatmaps in Figure 7 reveal that while cones of the same type were often very similar, they were not universally so - many produced statistically different mean color ratings. These effects were most clear in the green-red dimension of S20076 and to a lesser degree S20053 (Figure 7B,D). Overall, the yellow-blue dimension was more uniformly distributed across cone types (more yellow, less blue), owing to less frequent usage of these terms. The results from S20092, whose cone types were not known, are shown in Figure 7E,F. In general, a similar pattern emerged from this subject. Responses from cones were statistically distinguishable from about a third of the other cones. Together, this analysis confirms that many L- and M-cones elicited distinct hue sensations as hypothesized by Krauskopf and Srebro (1965). However, these results also indicate that variability between cones with the same photopigment also exists, which confirms the observations of Hofer et al. (2005b).

**Figure 7:**
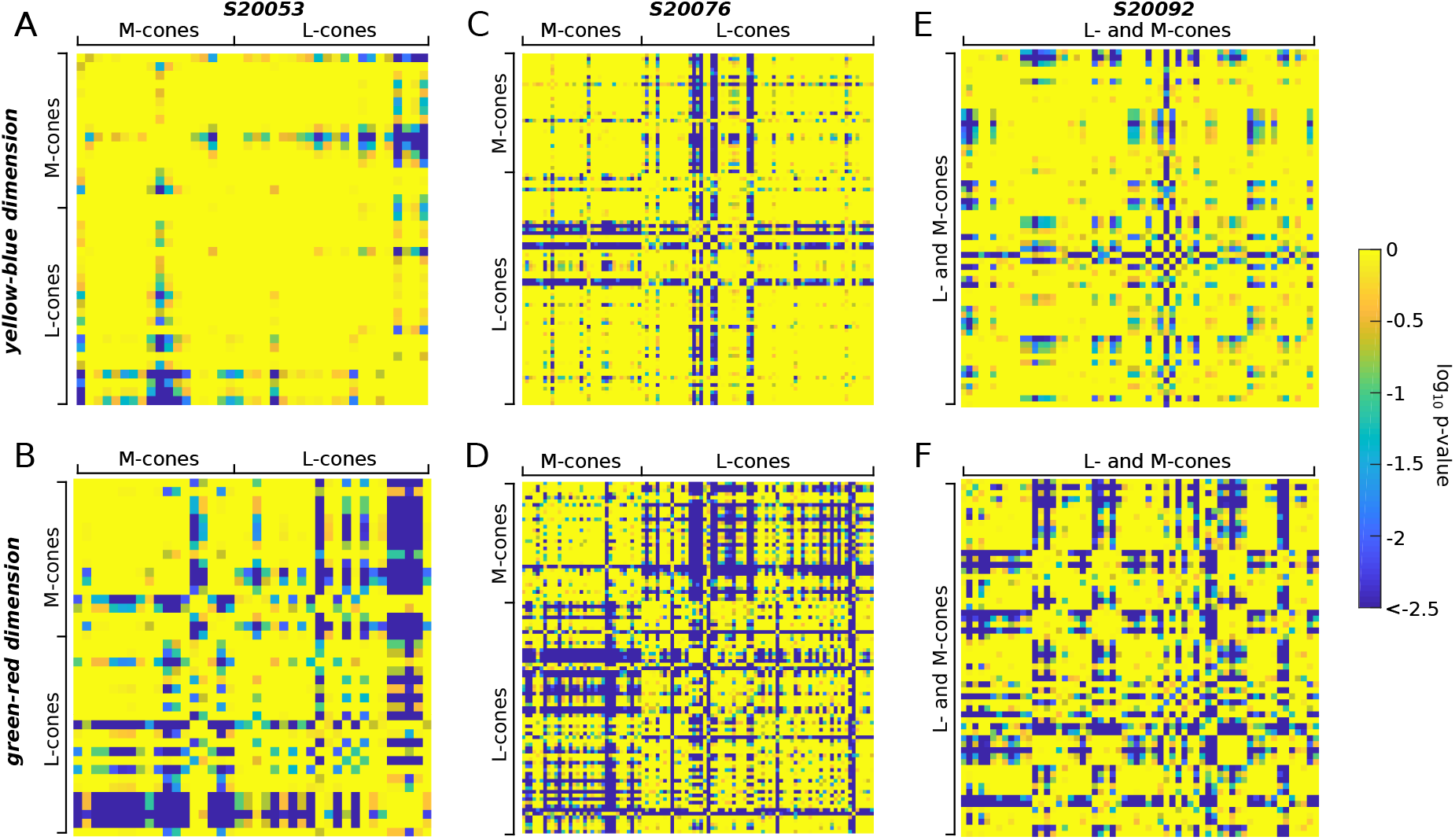
Cones with the same photopigment yield different sensations. An ANOVA revealed a main effect for cone targeted on hue/saturation scaling. The results of post-hoc tests (Tukey-Kramer) are shown. The mean response (UAD) measured for each cone was compared to all other cones. Comparisons were organized according to cone class. Heat maps represent the statistical significance of each comparison. Bluer colors indicate that the mean responses were statistically different (deep blue denotes *p* ≤ 10^−2.5^). Yellow-blue (**A, C, E**) and green-red (**B, D, F**) dimensions of UAD space were assessed separately. **A, B** S20053, **C, D** S20076, **E, F** S20092. The diagonal yellow line in each plot corresponds to the location where each cone was compared with itself. Each matrix is symmetric along the diagonal axis, *i.e*. the top and bottom triangles are mirror images.

### Hue percepts predict cone type

The tendency for cones to produce responses consistent with cone type encouraged us to ask how well cone type could be predicted from hue scaling. The results from S20053 and S20076 were collected into a single dataset and a support vector machine (SVM) was fit to the data. SVMs are a supervised approach to learning categorical labels. The SVM was given the position of each cone in UAD coordinates and its objectively classified (via densitometry) cone type and the algorithm learned a decision criterion. The learned boundary between L- and M-cones is shown by a solid diagonal line in Figure 8A. The trained SVM was then used to predict cone types based on the measured UAD position; a procedure we termed subjective classification. Following this procedure, it was found that subjective and objective classification agreed 79.0±3.5% (108/137) of the time, which was statistically higher than expected by chance (Cohen’s Kappa = 0.534, *p* < 0.001). Similar results were obtained when the SVM was trained on each subject’s data separately. Based on the robustness and accuracy of this procedure, the trained SVM was used to predict the data from S20092. The UAD data and SVM boundary are shown in Figure 8B and the spatial location of each subjectively classified cone is represented in Figure 8C. This procedure identified 33/60 L-cones. However, it is worth noting that S20053 and S20076 had very similar L:M cone ratios (near 2:1) and it is not known whether the SVM boundary would be influenced by this ratio.

**Figure 8:**
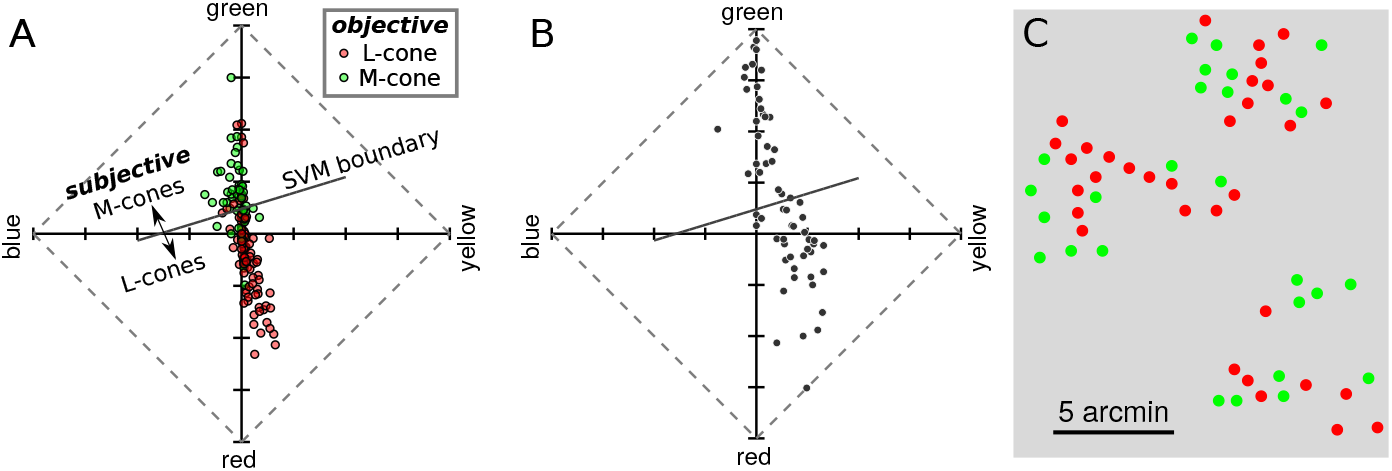
Hue and saturation reports predict cone type. **A** Data from S20053 and S20076 were collected into a UAD coordinates and fit with a SVM classifier. The classifier identified each cone as either L- or M-cone based on its mean UAD position. The learned decision boundary is shown by the solid diagonal line. This procedure is referred to as subjectively derived classification. For comparison, the objectively derived, densitometry based, classifications are represented by the color of each dot. **B** The results from S20092, whose cones have not been objectively classified, are re-plotted in an UAD and the SVM decision boundary learned in **A** is shown. Cones above the line were classified as M-cones and those below as L-cones. **C** Spatial location of each cone in the mosaic. The color of each dot indicates the subjectively inferred cone type (red=L, green=M).

### Possible influence of neighboring cones

Lastly, we explored the factors that potentially caused some cones to produce sensations that were inconsistent with their photopigment. One factor that could influence color sensations are other cones spatially proximal to the targeted cone. The local neighborhood of a targeted cone could influence perception in at least three ways: (1) differential baseline activity might either adapt or sensitize post-receptoral pathways (Schmidt et al., 2018; Tuten et al., 2017), (2) light from the flash might leak into neighboring cones and (3) neighboring cones may influence the hard-wired or learned hue sensation associated with each cone (Benson et al., 2014). In these experiments, the baseline activity of L- and M-cones was approximately equal and unlikely to have been a major factor. The latter two possibilities are worth considering in more detail.

The possibility that uncontrolled light leaking into neighboring cones had a significant influence on perception was tested first. The mean saturation of each cone was computed as a function of the number of non-like neighbors in its immediate surround. If the small proportion of light reaching neighboring cones were influencing hue and saturation judgments for single cone-targeted stimuli, one expectation is that saturation judgments should decrease as the number of non-like neighbors increases. The intuition being that if, for example, an L-cone with six surrounding M-cones was targeted, then light leaking into the neighboring M-cones would generate a less saturated (whiter) sensation (Krauskopf and Srebro, 1965). Alternatively, the local neighborhood may influence color judgment through prior expectation. Some models of small spot color appearance, *e.g*. Brainard et al. (2008), predict than an L-cone surrounded by six M-cones should produce a more saturated red sensation than a mixed surround, since post-receptoral pathways would carry a strong chromatically opponent signal in the former case. Figure 9 demonstrates that neither expectation was borne out: saturation ratings were not dependent upon the number of non-like neighbors. Similarly, the mean hue angle from each cone was not significantly influenced by the number of non-like neighbors (*p* > 0.05).

**Figure 9:**
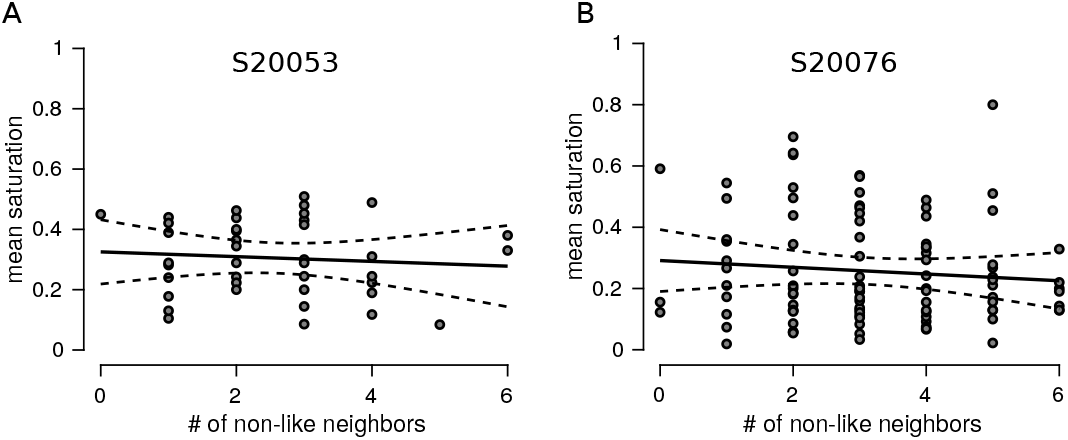
Influence of neighboring cones on saturation judgments. Each data points represents the mean saturation judgment as a function of number of non-like neighbors from a tested L- or M-cone in S20053 (**A**) and S20076 (**B**). Data includes all seen trials across all intensities. Solid black line indicates best fit regression and dotted lines are 95% confidence intervals. *p* > 0.05 for both correlation analyses.

## Discussion

The neural computation of color depends on both relative activity across the three cone types and previous experience (Brainard, 2015; Gegenfurtner, 2003). In the case of very small spots, like the ones used here, the visual system is unable to compare stimulus-driven activity across the cone classes and must rely on prior experience, or hard-wired connections, alone. Despite the unnatural nature of such a stimulus, subjects often see colored spots and we report here that hue sensations were predicted based on the photopigment of the cone probed (Figure 8). These results support the idea that, near the fovea, the visual system maintains information about the spectral class of each cone in its own receptor mosaic.

We found that the spectral identity of 79.0±3.5% of targeted cones could be correctly identified based on the color they generated (Figure 8) and as much as 50% of the variability in hue judgments could be attributed to the cone type targeted. These observations are remarkable for a few reasons. First, a light capture model indicated that the targeted cone did not absorb 100% of the stimulus light (Figure 1). Between 18 and 33% of absorbed light was captured by the nearest six cones. Theoretically, light leak into these nearby cones could have had a profound influence on the perceived color or saturation of stimuli. However, we did not find evidence in support of such an influence (Figure 9). This observation was consistent with the targeted cone acting as the dominant driver of hue reports. Second, the present subjective cone classification was at least as good as a recent study that was designed to classify cones psychophysically. Tuten et al. (2018), using adaptive optics, measured sensitivity to cone targeted flashes and compared those results to objectively identified cone classes. The authors used two chromatic backgrounds and stimulus wavelengths to selectively bias detection towards either L- or M-cones. With this procedure they found 77±2.9% of cones classes were correctly predicted. In the current paradigm, both baseline activity and sensitivity to the stimulus probe were equalized between L- and M-cones. Thus, at the level of the receptor mosaic, each stimulus flash produced an identical pattern of activity. Thirdly, the objective classification method itself has an error rate estimated to be between 3-5% (Sabesan et al., 2015), which sets the upper bound on how well any comparison can do. Despite all of these challenges and limitations, subjectively derived cone classification, based solely on hue and saturation scaling, agreed well with objective measurements. This evidence indicates that the visual system has prior information, either hard-wired during development or learned through experience, about each cone in its mosaic – a “lookup table” of sorts – that it leverages during the process of assigning a hue sensation.

The presence of a “lookup table”, which associates each receptor in the human cone mosaic with a type specific hue, may be surprising given the arrangement of the cone mosaic. There are no known molecular markers that differentiate L- from M-cones and the spatial topography of L- and M-cones near the fovea follows an approximately random distribution (Hofer et al., 2005a; Roorda and Williams, 1999; Roorda et al., 2001). During development, the expression of a single opsin gene in each cone is thought to arise through a stochastic process (Knoblauch et al., 2006; Neitz and Neitz, 2011). Furthermore, cones migrate during development towards the fovea (Hendrickson, 1994), which introduces additional spatial randomization. Thus, acquiring a spatially detailed representation of the cone mosaic is a challenge for the visual system. One possibility is that the brain learns a map of the location (Ahumada and Mulligan, 1990) and spectral identity of the cone mosaic through visual experience (Brainard, 2015; Benson et al., 2014). The intuition behind this idea is that during natural viewing the activity across a cone mosaic will be correlated in space, time and across spectral classes due to the statistics of natural scenes, eye movements and the action spectra of the photopigments. Benson et al. (2014) demonstrated that spatial and spectral correlations are theoretically sufficient to distinguish the spectral identity of cones in a mosaic. The parvocellular pathway is an obvious candidate for representing information at such fine spatial scales. Near the fovea, parvocellular (midget) ganglion cells receive input from a single cone (Kolb and Marshak, 2003; Dacey, 2000) and this private line is thought to be approximately preserved through at least the lateral geniculate nucleus (McMahon et al., 2000; Schein, 1988). In cortex, receptive fields of visual neurons increase in size (Dumoulin and Wandell, 2008; Felleman and Van Essen, 1991) and pool signals from more and more neurons. Therefore, information about cone type is unlikely to arise *de novo* at later centers.

Further insight into the neural pathways most likely involved in this task comes from the variability in color judgments when cones with the same photopigment were targeted (Figure 6&7). Classically, small spot experiments were interpreted in the tradition of Muller’s Law (Krauskopf, 1978; Krauskopf and Srebro, 1965; Krauskopf, 1964; Ingling et al., 1970; Otake and Cicerone, 2000; Hartridge, 1946). Within this framework a single neuron was thought to represent the presence or absence of a single variable, such as a red surface or an oriented bar – *i.e*. a labeled line (Muller, 1930). Accordingly, a higher firing rate was thought to indicate a redder stimulus, for example. Krauskopf and Srebro (1965) argued that the appearance of spots detected by a single cone would carry a single chromatic sensation: light absorbed by an L-cone would appear red, while an M-cone greenish-blue. Our results were only partially consistent with that prediction. Red/yellow was most frequently reported when an L-cone was targeted, while green/blue was used most often on M-cone trials (Figure 4&6). However, the present work, along with previous reports (Sabesan et al., 2016; Schmidt et al., 2018; Hofer et al., 2005b), demonstrated that cones within a spectral class often elicited different percepts (Figure 7) and the most common response was white. Together these observations contradict the strongest form of Krauskopf and Srebro’s hypothesis.

A single cone contributes to the receptive field of at least 20 classes of ganglion cells that each tile the retina (Masland, 2011; Dacey, 2004). For this study the two numerically dominant classes of ganglion cells – the parasol and midget classes – are most relevant for consideration. Both classes are known to be excited by single cone stimuli (Li et al., 2014; Freeman et al., 2015; Sincich et al., 2009) and project to the lateral geniculate nucleus (Dacey, 2004). Midget ganglion cells have been proposed as the retinal substrate for red-green chromatic sensations and high acuity vision (Lee, 2011; Dacey, 2000), while parasol ganglion cells constitute a channel carrying achromatic and motion information (Manookin et al., 2018; Schiller et al., 1990). Presumably each cone targeted in our study contributes to both midget and parasol channels. However, the relative strength of those connections may differ between cones (Li et al., 2014). Differences in connection strength could underly the within class variability in saturation ratings we observed. Were this the case, then altering the contrast (intensity) of the stimulus should favor one pathway over the other. At lower contrasts, parasol cells would be favored owing to their higher contrast response gain (Kaplan and Shapley, 1986; Shapley, 1990). At higher contrasts, the lower sensitivity midget (parvocellular) pathway will be more strongly recruited. Thus according to this idea, saturation should increase proportionally with intensity. However, we did not observe this phenomenon. Saturation judgments were relatively independent of stimulus intensity (Figure 4) and in two of the three subjects the relationship went in the opposite direction. These conclusions are further supported by previous work using small spots (Finkelstein and Hood, 1982, 1984; Finkelstein, 1988).

Alternatively, a population of neurons together may represent a property, such as color, as a probability distribution (Pouget et al., 2013). This could be accomplished if sensory input is encoded across a population of neurons with variable tuning (Ma et al., 2006; Finkelstein and Hood, 1984). Numerous authors have proposed that a representation of color space could be built in such a manner by cortical circuits that combine input from lower visual areas (Bohon et al., 2016; Kellner and Wachtler, 2013; Brainard et al., 2008; Emery et al., 2017; Finkelstein and Hood, 1984). For instance, midget ganglion cells multiplex chromatic and achromatic signals, an idea known as “double duty” (Rodieck, 1991; Dacey, 2000; Shapley, 1990). The relative strength of chromatic versus achromatic information varies between cells due to random wiring (Wool et al., 2018; Crook et al., 2011). Near the fovea, each midget cell would have an L- or M-cone in its center and variable cone weights in the surround, based on the cone types of its closest neighbors. Models explaining how randomly wired midget ganglion cells may sub-serve both chromatic and achromatic sensation have been described previously (Sabesan et al., 2016; Schmidt et al., 2014, 2016; De Valois and De Valois, 1993; Ingling Jr. and Martinez, 1983; Finkelstein, 1988; Finkelstein and Hood, 1984; Wool et al., 2018). Since each midget cell carries a unique chromatic/achromatic signature, the visual system could learn to associate different prior information with the output of each midget ganglion cell through experience (Benson et al., 2014). The color reported when a single cone is targeted with light may be a reflection of that prior experience. Finally, this theory makes the testable prediction that color reports in the peripheral retina, where the centers of midget-cell receptive fields pool signals from multiple cones (Dacey, 2000), will be less tightly predicted by the cone type stimulated.

In summary, using a middle wavelength small spot probe we found (1) brightness ratings and frequency of seeing depended upon stimulus intensity, but were not influenced by cone class. (2) Cones with the same photopigment often produced statistically different responses. (3) Despite this within cone class variability, hue responses varied so predictably between L- and M-cones that spectral identity was predicted from color reports with high accuracy. (4) Local neighborhoods had little, to no, influence on cone targeted sensations. These observations demonstrate that the visual system possesses a model of the world with enough precision to assign a meaningful hue label to spots of light that modulates the activity of only a single cone.

## Acknowledgments

We thank Professor Ramkumar Sabesan for providing helpful feedback on an earlier draft of this manuscript. We are grateful for technical assistance from Dr. Nicolas Bensaid, Dr. Francesco LaRocca and Pavan Tiruveedhula. This work was supported by grants from National Eye Institute National Institute of Health (R01EY023591, F32EY027637 and T32EY7043-38), the Minnie Flaura Turner Memorial Fund for Impaired Vision Research and the Michael G. Harris Ezell Fellowship.

